# RNAseq analysis reveals transcriptome changes associated with the progression of non-alcoholic liver disease in livers from *Efcab4b* knockout mice

**DOI:** 10.1101/2024.05.13.593681

**Authors:** Chew W. Cheng, Lucia Pedicini, Cintli Morales Alcala, Fenia Deligianni, Niamh Forde, Jessica Smith, Lynn McKeown

## Abstract

*EFCAB4B* is an evolutionarily conserved protein that encodes for the Rab GTPase Rab46, and the CRAC channel modulator, CRACR2A. Previous genome wide association studies have demonstrated the association of *EFCAB4B* variants in the progression of non-alcoholic liver disease (NAFLD). In this study we show that mice with global depletion of *Efcab4b^-/-^* have significantly larger livers than their wild-type (WT) counterparts. We performed RNA-sequencing (RNA-seq) analysis of liver tissues to investigate differential global gene expression among *Efcab4b^-/-^*and WT mice. Of the 69 differentially expressed genes (DEGs), analyses of biological processes found significant enrichment in liver and bile development, with 6 genes (*Pck1, Aacs, Onecut1, E2f8, Xbp1,* and *Hes1*) involved in both processes. Specific consideration of possible roles of DEGs or their products in NAFLD progression to (NASH) and hepatocarcinoma (HCC), demonstrated DEGs in the livers of *Efcab4b^-/-^* mice had roles in molecular pathways including lipid metabolism, inflammation, ER stress and fibrosis. The results in this study provide additional insights into molecular mechanisms responsible for increasing susceptibility of liver injuries associated with *EFCAB4B*.

## INTRODUCTION

Non-alcoholic fatty liver disease (NAFLD), a chronic liver condition characterised by excessive fat in the liver, is associated with insulin resistance, obesity and lipidaemia ^1^. In the UK, early-stage NAFLD may be present in 1 out of 3 people and is often asymptomatic. However, an increasing population of NAFLD patients will develop non-alcoholic steatohepatitis (NASH: defined as the presence of more than 5% hepatic steatosis and inflammation with hepatocyte injury^2^). The persistence of inflammation can progress NASH to hepatic cirrhosis and hepatocellular carcinoma (HCC) ^3^ ^4^ ^5^. Whilst NAFLD is the leading cause of liver fibrosis and HCC worldwide ^6^, it is also associated with the development of non-liver adverse outcomes such as cardiovascular diseases ^7^ ^8^ ^9^ and type 2 diabetes mellitus ^10^. In addition to the effect on wellbeing and quality of life, the total economic costs of diagnosed NASH in the UK is estimated to be £2.3 to £4.2 billion ^11^.

The progression of NAFLD to NASH is multifactorial, however, the underlying mechanisms are not well understood. Whilst a population of patients develop NASH some, with the same co-morbidities and risk factors, just have a fatty liver. This variation in disease progression suggests genetic predispositions and, indeed, studies have shown that genetic factors play a role in NAFLD pathogenesis ^12^ ^13^. Although a polygenic disease, common genetic variants have been consistently associated with increased risk of steatosis, particularly small nucleotide polymorphisms (SNPs) in Patatin-like phospholipase domain-containing protein 3 (*PN<u>P</u>LA3*), Lysophospholipase-like 1 (*LYPLAL1*), protein phosphatase 1 regulatory subunit 3B (*PPP1R3B*), Neurocan (*NCAN*) and glucokinase regulator (*GCKR*) ^14^ ^15^ ^16^. However, an accumulation of fat is not the only risk factor for promoting the transition from non-alcoholic fatty liver to NASH, a central component is persistent inflammation ^4,5^. Identifying the triggers of inflammation remains a major issue in the field. Recently a pilot GWAS in patients with NAFLD looked at genetic variants significantly associated with hepatic histology. Here they identified rs887304 on chromosome 12 in EF-Hand calcium Binding Protein 4B (*EFCAB4B*) (also known as Ca^2+^ Release-Activated Channel Regulator 2A; CRACR2A) to be associated with lobular inflammation ^17^. To elucidate a genetic predisposition to the pathogenesis and progression of NASH, Grove *et al*. explored multiple risk-associated alleles present in monozygotic twins that both developed NASH cirrhosis and revealed both patients were heterozygous for six SNPs, including rs887304 in *EFCAB4B* ^18^. Moreover, rs887304 was one of 19 SNPs shown to be significantly associated with NAFLD in a pilot study in an Indian population ^19^.

*EFCAB4B* encodes for two functional proteins: CRACR2A (CRACR2A-S, CRACR2A-201) ^20^ and Rab46 (CRACR2A-L, CRACR2A-203) ^21^. rs887304 is located at the 3’ UTR of CRACR2A but is intronic in the Rab46 coding region and could potentially regulate isoform expression (https://www.ensembl.org/Homo_sapiens/Gene/Variation_Gene/Table?db=core;g=ENSG00000130038;r=12:3606633-3764819). CRACR2A has been shown to be a regulator of store-operated calcium in T-cells ^22^ whilst Rab46 regulates the trafficking of unique granules in endothelial cells ^23^ and differential secretion in mast cells ^24^. In endothelial cells, Rab46 acts as a brake to prevent the secretion of pre-stored pro-inflammatory components in response to non-thrombotic stimuli. The depletion of Rab46 in endothelial cells could therefore lead to the increased inflammation necessary for NAFLD disease progression, however a patient with biallelic mutations in EFCAB4B displayed immunodeficiency due to a loss of function in T-cells ^25^. Since SNPs in *EFCAB4B* ^26^ and the presence of NAFLD ^27^ have been associated with severe and critical responses to COVID-19, outcomes that are associated with an extreme inflammatory response, it is important to understand the contribution of CRACR2A and Rab46 to liver function. Here we utilized an *Efcab4b* global knockout mouse to explore the impact of CRACR2A and Rab46 depletion on liver and to identify some potential pathways that contribute to inflammation/progression of NAFLD.

## METHODS

### Study design

#### Mice

Murine work was carried out in accordance with The Animals (Scientific Procedures) Act 1986 (Amended 2012). Mice were kept in the University of Leeds animal facility under standard conditions, including a 12-hour sleep/wake cycle, with access to water and chow diet ad libitum. Experiments were conducted under Home Office Project License P606230FB. All studies were approved by the University of Leeds Ethics Committee. Male and female animals were ear notched at weaning, with these samples subsequently used for genotyping.

#### Generation of *Efcab4b^-/-^* mice

The *Efcab4b* knockout mouse strain was created from ES cell clone 15424A-C4, generated by Regeneron Pharmaceuticals, Inc. and litters obtained from the KOMP Repository (www.komp.org). The homozygous C57BL/6N-CRACR2At^m1.1^(KOMP)^Vlcg^ and WT C57BL/6N strains were achieved by heterozygous matings. Mice were depleted of the gene *Efcab4b* which prevents the expression of both CRACR2A and Rab46 protein. Genotyping was performed and homozygous and WT mice were selected for breeding. Mouse genotyping was performed by taking ear notches at weaning age. The samples were sent for automated genotype PCR service (Transnetyx, Cordova, TN, USA; see primer Table 1).

**Table 1.**
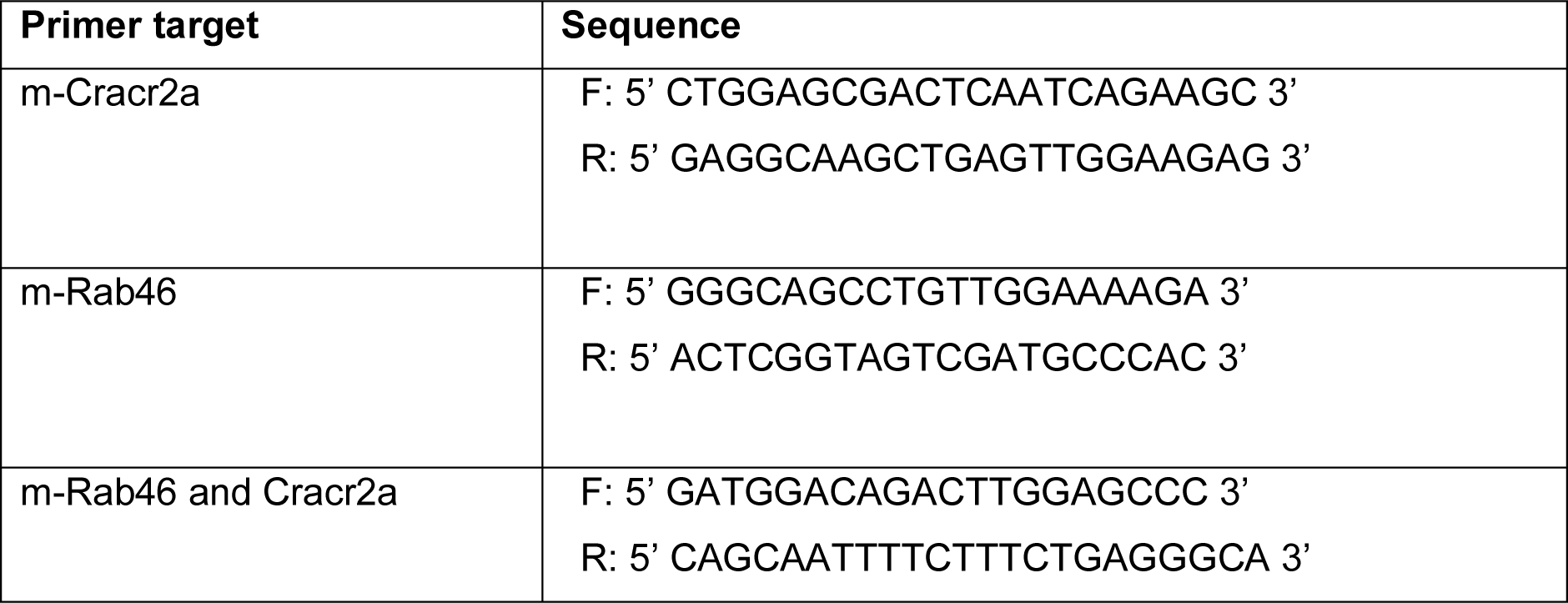
PCR primers.

### Sampling

For histology, homozygous KO (*Efcab4b*^-/-)^ and WT (*Efcab4b*^+/+^) mice were weighed at 8 weeks and 12 weeks after which they were schedule 1 and organs were collected. The dissected organs were weighed prior to being snap frozen in liquid nitrogen and stored at in paraformaldehyde for histological examination. Organ-to-body weight ratios were calculated normalising the weight of each organ with its respective body weight.

### Histology

Liver samples were fixed in paraformaldehyde overnight and then kept in 70% ethanol until further analysis. Liver tissues from female *Efcab4b^-/-^* and WT mice were processed at the St. James’s hospital histology facility (Leeds). Histological examination of liver tissue was performed from the right lobe of the liver using Haematoxylin and Eosin (H&E) staining to reveal the main hepatic features. H&E staining was performed by the histology department at St. James’s hospital and imaged slices were sent and opened with Aperio ImageScope for analysis.

### RNA isolation

Liver tissue from 3x *Efcab4b^-/-^* male mice 11 weeks and 3x *Efcab4b^+/+^* mice 11 weeks were placed in an RNA/DNAse free tube with a metal lysis bead with 1 ml Trizol Reagent. Tubes placed in a tissue lyser were agitated at 26 Hz for 3 x 1 min. The resultant liquid was transferred to 1.5 ml centrifuge tubes, 200 μl of phenol-chloroform was added for 3 mins at room temperature, then spun at 12000g (4°C) for 15 mins. The supernatant (top aqueous phase) was transferred to new tubes and 500 μl of isopropanol added to each tube and incubated for 10 mins. The samples were spun for 15 mins at 12000g (4°C), and the supernatant discarded. The pellets were re-suspended in 1 ml of 75% ethanol (in dH2O) and centrifuged at 8000g (4°C) for 5 mins. The supernatant was removed carefully, the pellet dried and then 20-50 μl of RNA-free dH2O was added to each sample and stored at -80°C. RNA was quantified using a NanoDrop®.

### RNA sequencing and bioinformatics analysis

RNA extracted from the three samples of each of the two conditions; WT and *Efcab4b*^-/-^, were sequenced by Novogene, Cambridge, United Kingdom. The TruSeq Stranded mRNA kit (Illumina) was used for library preparation, following the manufacturer’s recommended protocol. The libraries were sequenced on the HiSeq 4000 platform with 150 bp paired-end strategy. On average, 84.59 M high-quality reads were generated from the RNA sequencing project. The 24 raw reads were uploaded to the ENA-EMBL-EBI database under the accession number E-MTAB-13110.

STAR aligner^28^ was used to map the raw reads to the *mouse* reference genome (Ensembl GRCm39)^29,30^. The gene expression levels were quantified by featureCounts^28,31^ and normalised by DESeq2^32^ using the negative binomial model. Differentially expressed genes were defined based on adjusted p-value < 0.05 (FDR). For pathway enrichment analyses, Kyoto Encyclopedia of Genes and Genomes (KEGG) and REACTOME databases were searched via clusterProfiler and ReactomePA, respectively, to predict potential enriched pathways. The significant terms were selected based on adjusted p-value < 0.05. The principal component analysis plot, volcano plot and dot plots were generated using the GGPlot2 function in R.

Data availability via ArrayExpress https://www.ebi.ac.uk/biostudies/arrayexpress/studies/E-MTAB-13110.

## RESULTS

### Animal model validation

The *Efcab4b^-/-^* mouse strain did not show any viability or fertility issues with regards to fertilisation and litter size. Mice with disrupted CRACR2A/ Rab46 expression (*Efcab4b^-/-^* mice) and respective WT controls were sacrificed at 12 weeks and gene knockout was validated from the stated tissues (Figure 1). Quantitative RT-PCR analysis of mRNA abundance in the liver, lung, spleen and heart from *Efcab4b^-/-^* and WT mice using three sets of primers (see Table 1) specific for CRACR2A, Rab46 or both isoforms showed reduced mRNA expression of both the isoforms in all the selected organs, confirming global knockout of murine *Efcab4b* (Figure 1A). Moreover, CRACR2A and Rab46 knockout was confirmed at the protein level in mouse liver endothelial cells (mLECs) (Figure 1B). Immunoblotting using an antibody that recognises both isoforms exhibited specific staining at the expected molecular weight (95 kDa) corresponding to Rab46 in control mLECs and reduced intensity in *Efcab4b^-/-^* mLECs. These results confirm Rab46 deletion in mLECs. CRACR2A is not expressed in mouse or human endothelial cells.

**Figure 1.**
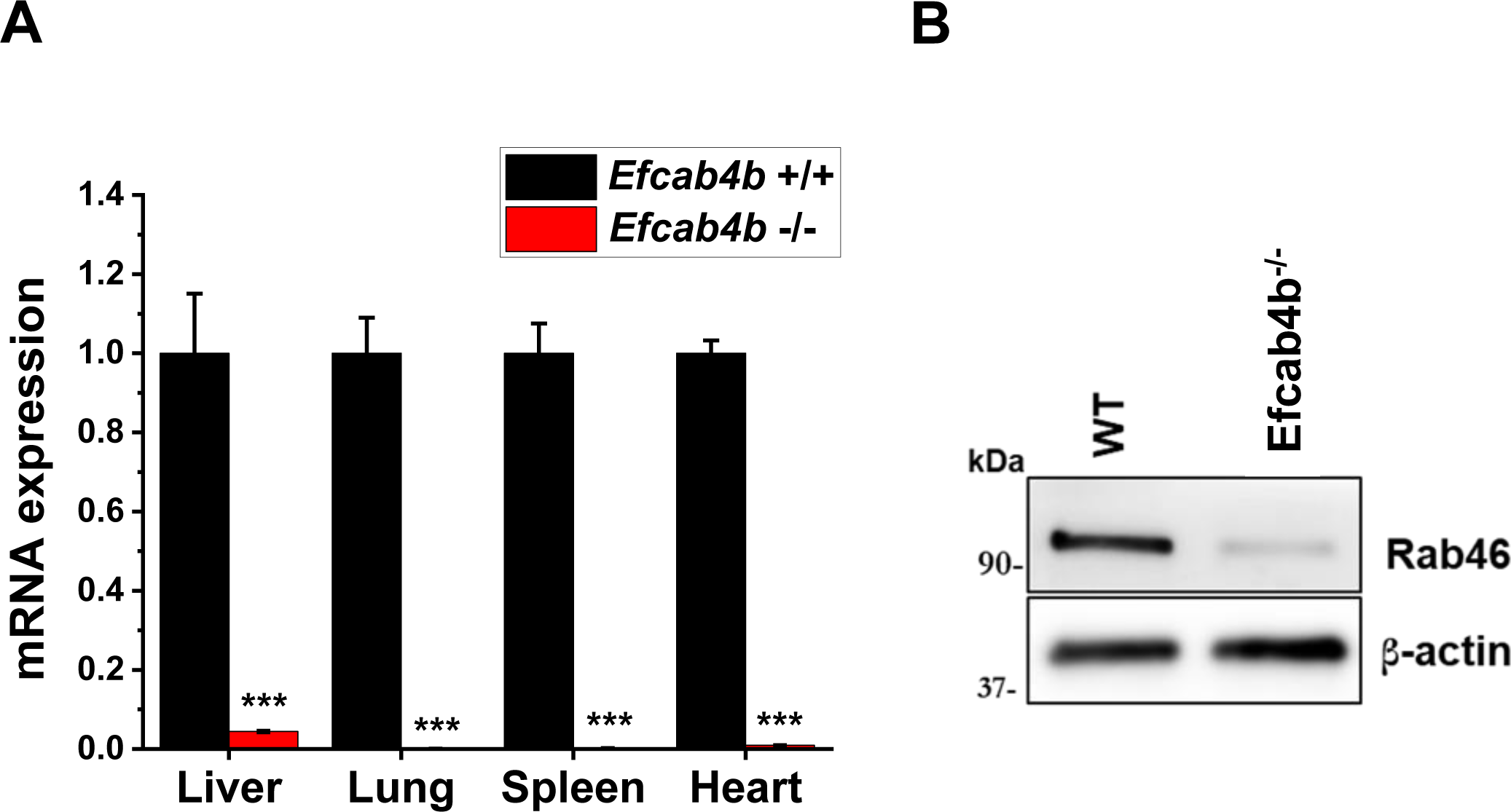
Validation of *Efcab4b* gene knockout in mice. (A) qPCR ΔCt analysis of *Efcab4b* expression relative to housekeeping genes and normalized to wild-type (WT) control in the liver, lungs, spleen and heart of *Efcab4b^-/-^* mice and WT control mice. The abundance of mRNA encoding for both the CRACR2A and Rab46 protein isoforms is significantly decreased in all the tissues. n=7 *** p<0.001. (B) Representative western blot to show Rab46 protein at the expected band size of ∼90kDa in lysates from LECs extracted from WT and *Efcab4b^-/-^* mice, using an antibody that recognises both CRACR2A and Rab46.

*Efcab4b^-/-^* mice appeared superficially normal in physical appearance compared to WT mice with no difference in life span up to 12 weeks between genotypes. Mice fed with standard chow diet had their body weight assessed at 8 and 12 weeks (Figure 2A). No difference was seen in body weight at the considered time points between WT (*Efcab4b^+/+^*) and *Efcab4b^-/-^* mice. A significant difference was observed in liver weight, where *Efcab4b^-/-^* mice have increased liver mass compared to WT mice (Figure 2B). A small but significant increase in the visceral fat was also observed in *Efcab4b^-/-^* mice.

**Figure 2.**
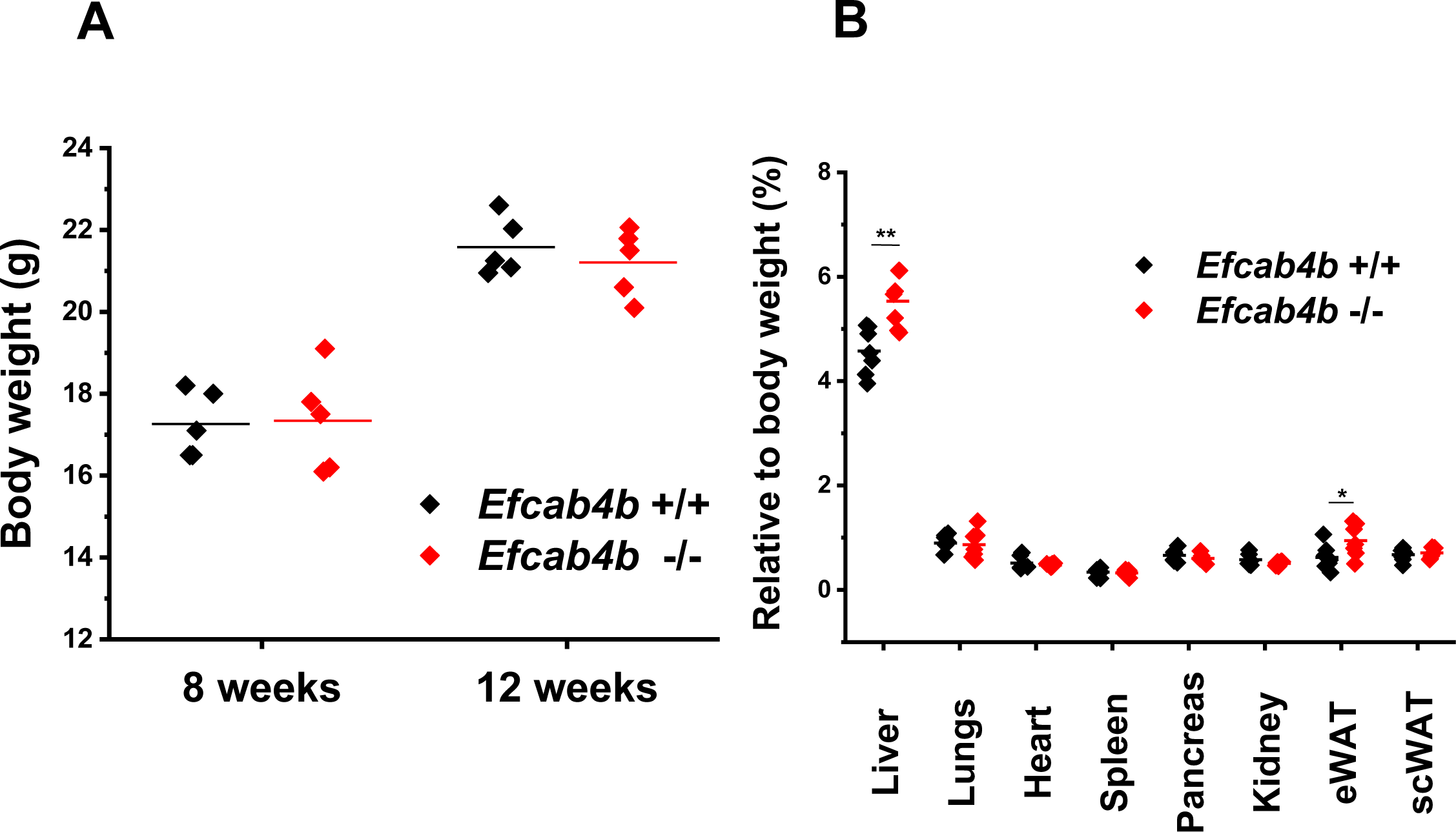
Effect of *Efcab4b* gene knockout on body and organ weights. A) Body weight at the age of 8 and 12 weeks of female control (WT: n = 5) and female *Efcab4b^-/-^* mice (n = 5). Scatter plot shows body weight (g) of each mouse and straight line indicates the group mean. No significant difference is observed at these time points between WT (black diamonds) and knockout mice (red diamonds). B) Male and female mice were sacrificed at the age of 12 weeks and the main organs were harvested and weighted. Scatter plot shows data distribution of each organ from control and *Efcab4b^-/-^* mice. Straight line indicates the mean value. Liver weight and eWAT weight is increased in *Efcab4b^-/-^* mice compared to control (WT). n=5 **p<0,01 *p<0,05 from Student t-test.

Histological analysis of livers from the WT mice indicated a normal liver lobular architecture with central vein and surrounding hepatocytes, sinusoids and nucleus (Figure 3). *Efcab4b^-/-^* mice show normal liver morphology, however hepatic mononuclear cell infiltration, congestion of portal vein and blood sinusoids appear to be a common feature. Accumulation of fat droplets was not detected in any of the analysed tissues section. The presence of focal periportal immune cell infiltration, with enlargement of the portal tract may suggest an early inflammatory condition due to gene deletion (Figure 3).

**Figure 3.**
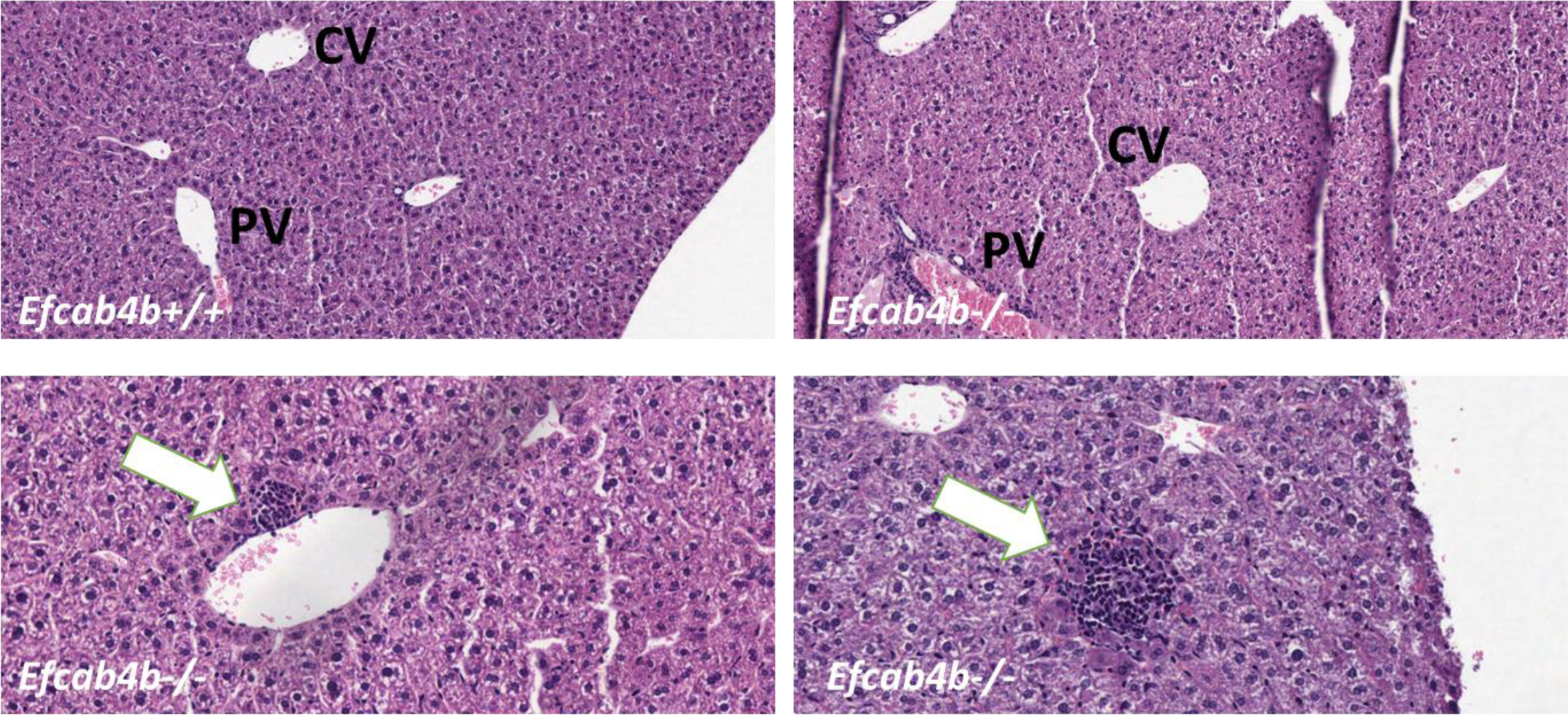
Histological examination of liver of *Efcab4b* KO mice compared to control (WT). Mice were sacrificed at the age of 12 weeks and the liver was harvested and weighed. Liver tissues were examined by H&E staining. Liver from WT *Efcab4b^+/+^* control group revealing normal morphology with central vein (CV) and normal triad structure with portal vein (PV); Liver from *Efcab4b^-/-^*mice reveals multiple focal inflammation sites with immune cell infiltration (white arrows) and portal vein and sinusoids congestion.

### Transcriptional changes suggest *Efcab4b* transcripts contribute to liver physiology

Having defined some of the morphological changes in the liver of *Efcab4b^-/-^* mice we undertook bulk RNA-seq analysis of the livers extracted from WT control and *Efcab4b^-/-^* mice. We compared the gene expression levels and associated functions of the genes differentially expressed between the two groups, to understand the influence of CRACR2A and Rab46 protein deficiency on mouse liver function.

A principal component analysis (PCA) demonstrated the source of greatest variation in transcriptional response of liver tissue (Figure 4) was source of tissue The analysis revealed that PC1 accounted for 33% of the variance, indicating a substantial contribution to the overall genetic variation between the samples. Similarly, PC2 explained 30% of the variance, further capturing significant differences in the genetic profiles. By comparing three knockout samples to three control samples, distinct patterns and clustering were observed in the PCA plot, suggesting notable genetic distinctions between the two groups.

**Figure 4.**
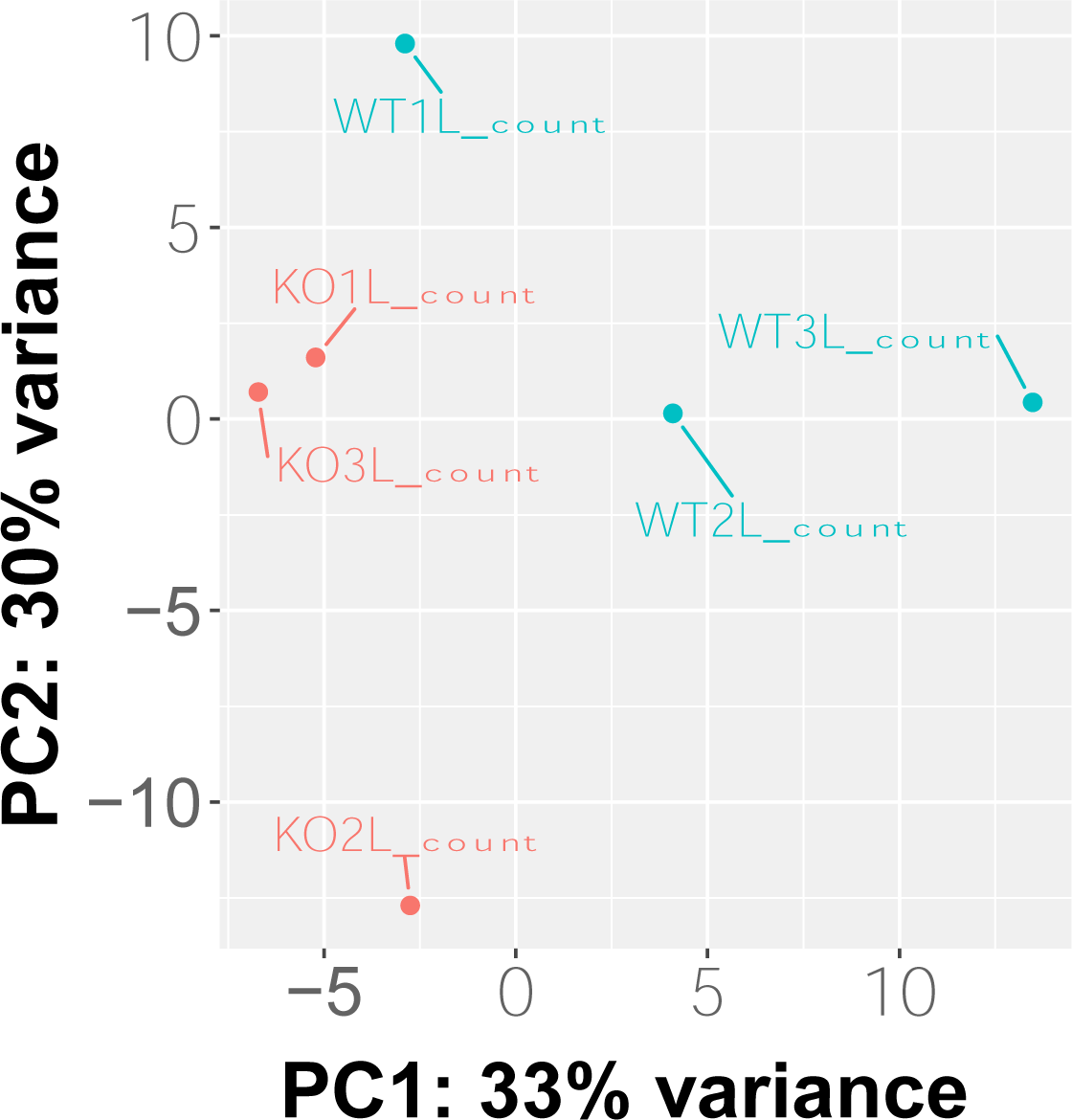
PCA plot showing segregation of samples based on genome-wide expression profiles. Each dot represents one sample, blue colour dots representing wild-type samples and red colour dots representing knockout samples.

Our study focused on comparing *Efcab4b^-/-^* and WT models, revealing a total of 69 DEGs (Figure 5). Among these DEGs, 30 were found to be upregulated while 39 were downregulated in the KO model. The full name of the top 30 differentially regulated genes in the *Efcab4b^-/-^* livers compared to *Efcab4^+/+^* (plus Log2 fold changes values and FDR) and their functional characteristics are shown in Table 2. Of those 30 upregulated and 39 downregulated DEGs, a simple literature search demonstrated 16 and 16 respectively were involved in molecular pathways that when dysregulated play a role in NAFLD and the progression to NASH and HCC (Tables 3 and 4).

**Figure 5.**
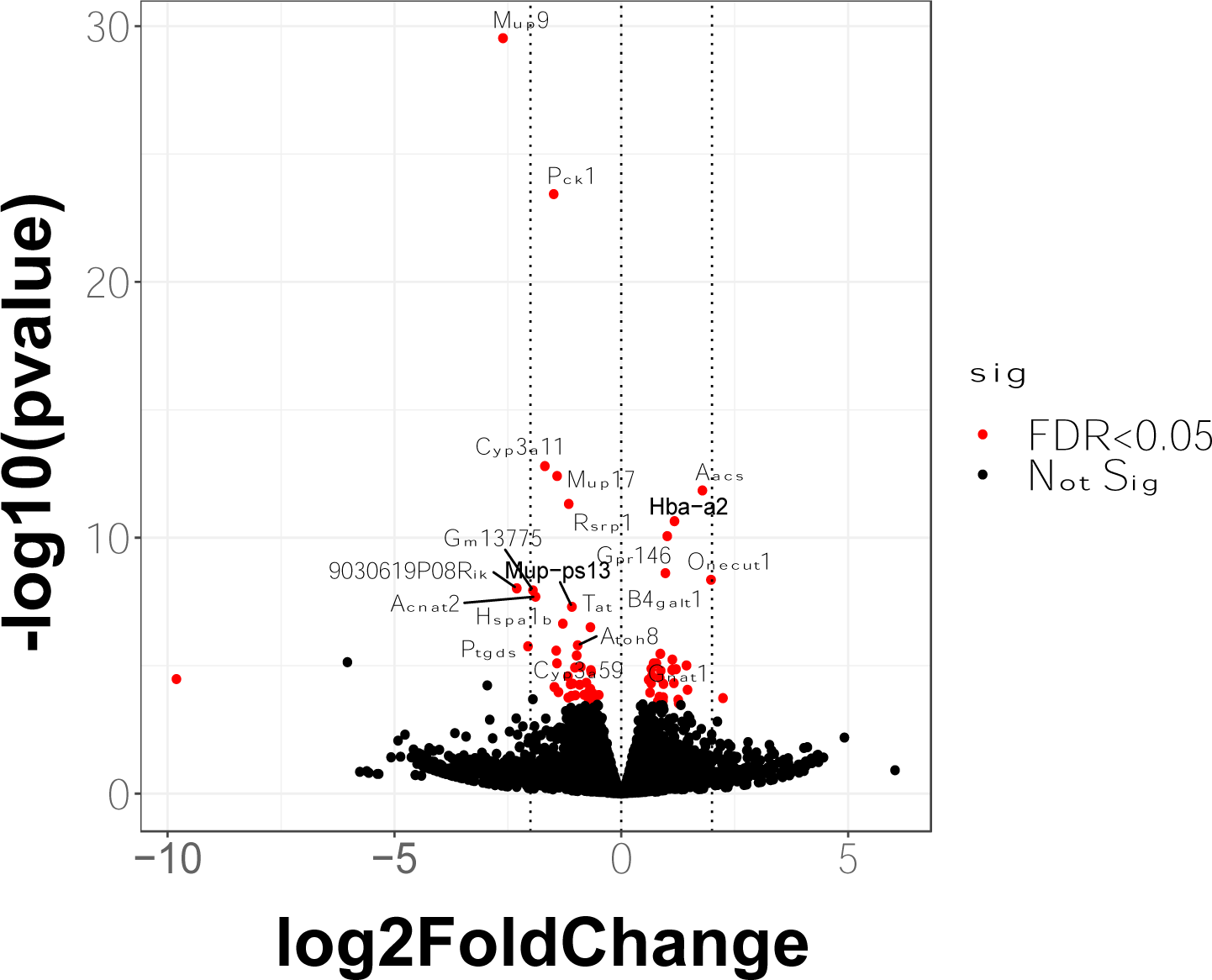
Volcano plot showing differentially expressed genes. based on adjusted P-value<0.05. There are 69 differentially expressed genes (DEGs) detected comparing between knocked-out and wild-type. Of those 69 DEGs, 30 DEGs are upregulated and 39 DEGs are downregulated in knocked-out model.

**Table 2.** The top 30 significantly differentially expressed genes (DEGs) based on adjusted p-value.

|  | GENE SYMBOL | LOG<sub>2</sub>FC | PADJ |
| --- | --- | --- | --- |
| 1 | Mup9 | -2.6075 | 3.1900E-26 |
| 2 | Pck1 | -1.4881 | 1.9800E-20 |
| 3 | Cyp3a11 | -1.6837 | 5.7200E-10 |
| 4 | Mup17 | -1.4155 | 1.0400E-09 |
| 5 | Aacs | 1.7884 | 3.0600E-09 |
| 6 | Rsrp1 | -1.1571 | 8.5500E-09 |
| 7 | Hba-a2 | 1.1723 | 3.4900E-08 |
| 8 | Gpr146 | 1.0139 | 1.1700E-07 |
| 9 | B4galt1 | 0.9729 | 2.9000E-06 |
| 10 | Onecut1 | 1.9776 | 4.7300E-06 |
| 11 | 9030619P08Rik | -2.3037 | 9.3100E-06 |
| 12 | Gm13775 | -1.9459 | 1.0300E-05 |
| 13 | Acnat2 | -1.8897 | 1.6800E-05 |
| 14 | Mup-ps13 | -1.0885 | 3.8500E-05 |
| 15 | Hspa1b | -1.2885 | 0.0002 |
| 16 | Tat | -0.6794 | 0.0002 |
| 17 | Atoh8 | -0.9598 | 0.0010 |
| 18 | Ptgds | -2.0571 | 0.0011 |
| 19 | Cyp3a59 | -1.4336 | 0.0015 |
| 20 | Gnat1 | 0.8614 | 0.0019 |
| 21 | Ces2c | -0.9825 | 0.0021 |
| 22 | Loxl4 | 1.1242 | 0.0028 |
| 23 | Serpina4-ps1 | 0.7744 | 0.0035 |
| 24 | Mvk | 0.7144 | 0.0035 |
| 25 | Gm49395 | -1.4180 | 0.0035 |
| 26 | E2f8 | 1.4379 | 0.0040 |
| 27 | Mup7 | -0.8927 | 0.0045 |
| 28 | Dio1 | -1.0201 | 0.0046 |
| 29 | Xbp1 | 0.6627 | 0.0049 |
| 30 | Fgfr1 | 1.2086 | 0.0049 |

**Table 3.** Upregulated genes with roles in pathways associated with NAFLD progression.

| GENE SYMBOL | LOG<sub>2</sub>FC | PADJ | LIVER DISEASE ROLE | GENE DESCRIPTION | EXPRESSION IN LIVER DISEASE | REF |
| --- | --- | --- | --- | --- | --- | --- |
| Socs2 | 2.24 | 0.0323 | NASH | Suppressor of cytokine signaling 2 | Down | 44 46 45 |
| Onecut1 | 1.98 | 0 | HCC | One cut domain, family member 1 | Down | 60 |
| E2f8 | 1.44 | 0.004 | HCC | E2F transcription factor 8 | Up | 61 |
| Fgfr1 | 1.21 | 0.0049 | Steatosis/HCC | Fibroblast growth factor receptor 1 | Up | 62 63 |
| Hes1 | 1.16 | 0.0123 | HCC | Hes family bHLH transcription factor 1 | Up | 64 |
| Sik1 | 1.13 | 0.0049 | steatosis | Salt inducible kinase 1 | Down | 65 |
| Loxl4 | 1.12 | 0.0028 | HCC | Lysyl oxidase-like 4 | Up | 66 |
| Gpr146 | 1.01 | 0 | Steatosis | G protein-coupled receptor 146 | Down | 33 |
| B4galt1 | 0.97 | 0 | HCC | UDP-Gal:betaGlcNAc beta 1,4-galactosyltransferase, polypeptide 1 | Down | 59 |
| Vgll4 | 0.92 | 0.0311 | HCC | Vestigial like family member 4 | Up | 67 |
| Idi1 | 0.92 | 0.0374 | Steatosis | Isopentenyl-diphosphate delta isomerase | Up | 38 |
| Prss8 | 0.87 | 0.005 | Steatosis | Protease, serine 8 (prostasin) | Down | 68 |
| Gnat1 | 0.86 | 0.0019 | HCC | Guanine nucleotide binding protein, alpha transducing 1 | Down | 69 |
| Odc1 | 0.84 | 0.03 | HCC | Ornithine decarboxylase, structural 1 | Up | 70 |
| Pmvk | 0.67 | 0.0076 | Steatosis | Phosphomevalonate kinase | Up | 39 |
| Xbp1 | 0.66 | 0.0049 | Steatosis | X-box binding protein 1 | Up | 40 41 |

*HCC: Hepatocarcinoma*

**Table 4.**
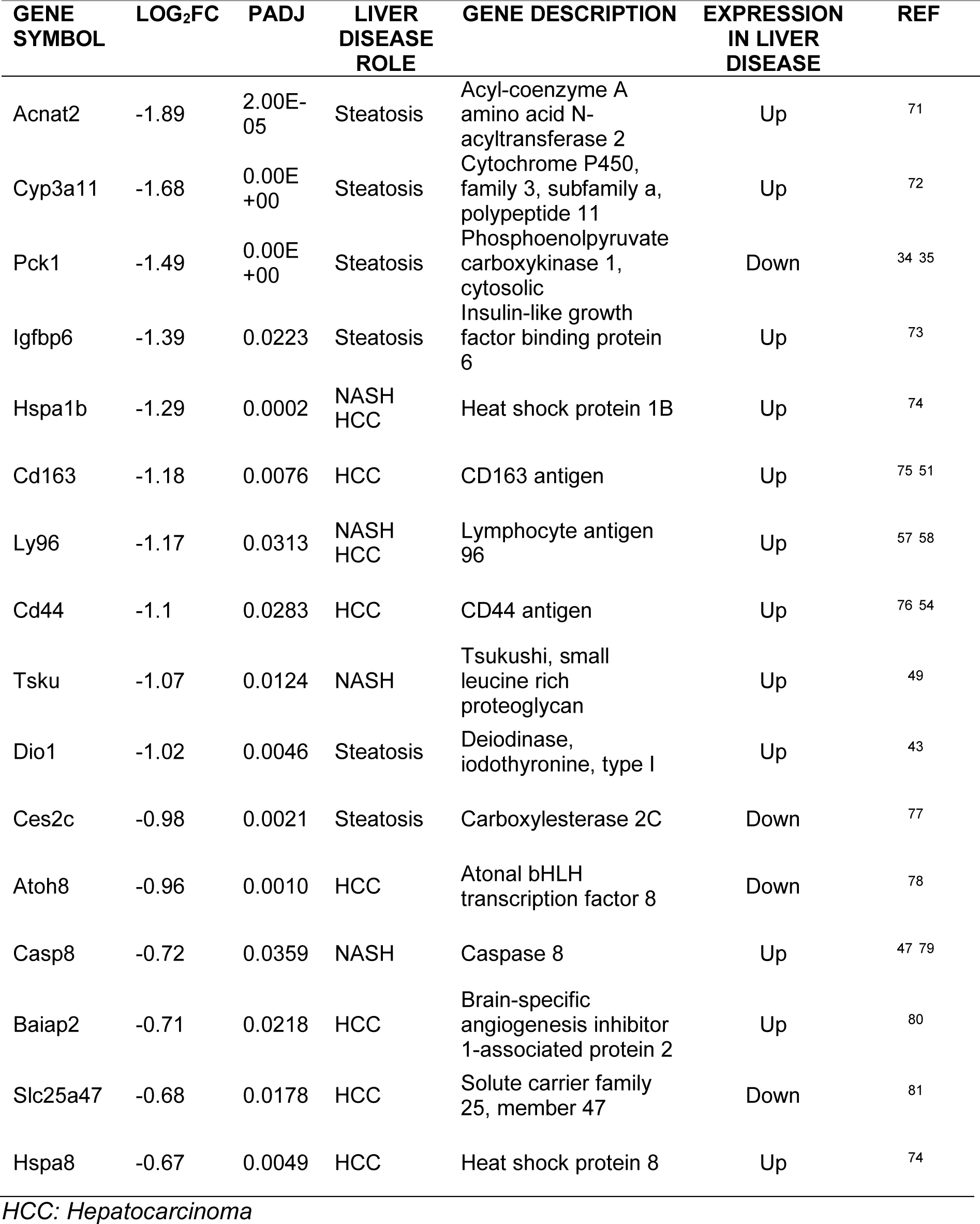
Downregulated genes with roles in pathways associated with NAFLD progression.

### Bioinformatic pathway analyses

Enrichment analysis was performed to gain insights into the functional significance of different Gene Ontology (GO) terms. Using all differentially expressed genes, we identified 55 significant terms based on adjusted P-value 0.05 (Figure 6A). The analysis revealed several enriched biological processes (BP) and molecular functions (MF) associated with the studied genes. In terms of biological processes, the study found significant enrichment in liver development (GO:0001889) and hepaticobiliary system development (GO:0061008), with 6 common genes (*Pck1, Aacs, Onecut1, E2f8, Xbp1*, and *Hes1*) being involved in both processes. Gland development (GO:0048732) was also enriched, with 9 genes (*Pck1, Aacs*, *Onecut1, E2f8, Xbp1, Fgfr1, Hes1, Cd44*, and *Socs2*) associated with this process. Furthermore, the enrichment analysis highlighted molecular functions such as estrogen 16-alpha-hydroxylase activity (GO:0101020) and retinoic acid 4-hydroxylase activity (GO:0008401), both mediated by the genes *Cyp3a11, Cyp3a59*, and *Cyp3a25*. Other enriched molecular functions included protein folding chaperone activity (GO:0044183), histone deacetylase binding (GO:0042826), and magnesium ion binding (GO:0000287).

**Figure 6.**
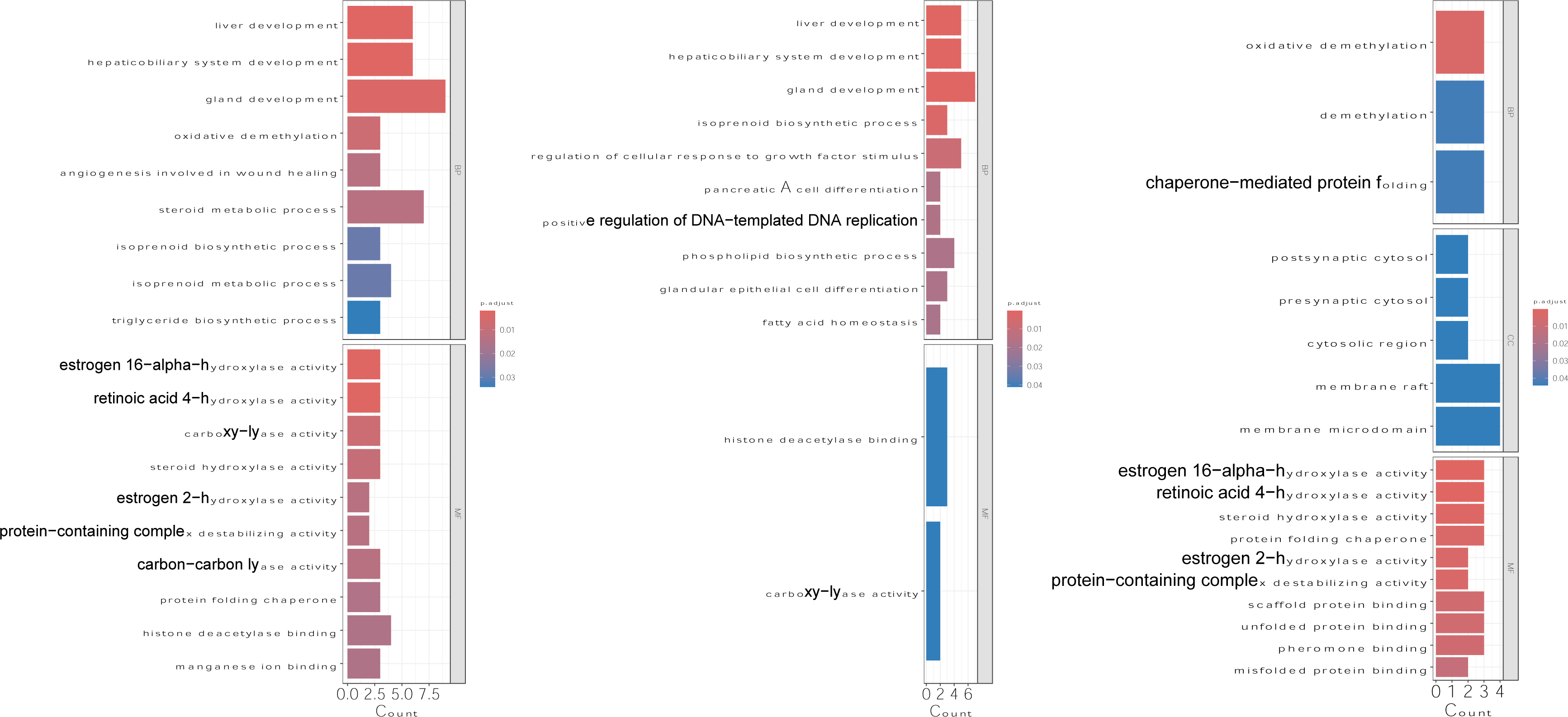
Enrichment analysis using differentially expressed genes. (A) Enrichment analysis of all DEGS; (B) enrichment analysis of upregulated DEGs; (C) enrichment analysis of downregulated DEGs. BP: biological processes, MF: molecular functions, and CC: cellular components.

Using only the upregulated differentially expressed genes. The analysis revealed 36 significantly enriched GO terms related to various biological processes and molecular functions (Figure 6B). In terms of biological processes, liver development (GO:0001889) and hepaticobiliary system development (GO:0061008) were found to be significantly enriched, Additionally, gland development (GO:0048732) showed enrichment. These findings suggest that the upregulated genes are associated with the development and function of liver and glandular tissues. Furthermore, the analysis revealed enrichment of genes involved in isoprenoid biosynthetic process (GO:0008299) and phospholipid biosynthetic process (GO:0008654). Genes related to monosaccharide metabolic process (GO:0005996) and hexose metabolic process (GO:0019318) were also enriched, indicating the involvement of upregulated genes in carbohydrate metabolism. Moreover, several enriched GO terms related to cellular signalling and regulation were identified. These included the regulation of cellular response to growth factor stimulus (GO:0090287) and the regulation of transforming growth factor beta receptor signalling pathway (GO:0017015). These findings suggest that the upregulated genes may play a role in modulating cellular responses to growth factors and signalling pathways. In terms of molecular functions, histone deacetylase binding (GO:0042826) and carboxy-lyase activity (GO:0016831) were significantly enriched. These findings indicate that the upregulated genes may be involved in epigenetic regulation and catalytic activities.

The enrichment analysis of the downregulated genes revealed 24 significant GO terms (Figure 6C). In terms of biological processes, the downregulated genes were enriched in processes such as oxidative demethylation (GO:0070989) and demethylation (GO:0070988), suggesting potential involvement in the regulation of methylation processes. Additionally, the downregulated genes were associated with chaperone-mediated protein folding (GO:0061077), indicating a disruption in protein folding mechanisms. In terms of cellular components, the downregulated genes were enriched in regions such as postsynaptic cytosol (GO:0099524), presynaptic cytosol (GO:0099523), cytosolic region (GO:0099522), membrane raft (GO:0045121), and membrane microdomain (GO:0098857), suggesting potential alterations in synaptic and membrane organization. The molecular function analysis revealed enrichment of activities such as estrogen 16-alpha-hydroxylase activity (GO:0101020), retinoic acid 4-hydroxylase activity (GO:0008401), steroid hydroxylase activity (GO:0008395), protein folding chaperone (GO:0044183), and scaffold protein binding (GO:0097110).

We performed a pathway enrichment analysis of all differentially expressed genes using the REACTOME database. The analysis revealed enrichment of genes involved in Xenobiotics (R-MMU-211981) metabolism, with the downregulated genes including *Cyp3a11, Cyp3a59,* and *Cyp3a25*. Additionally, the cholesterol biosynthesis (R-MMU-191273) pathway showed enrichment, with *Mvk, Pmvk,* and *Idi1*.

REACTOME pathway analysis was performed using upregulated genes. This analysis revealed two significant pathways were enriched by the differentially expressed genes. The first pathway, Cholesterol biosynthesis (R-MMU-191273), involves the synthesis of cholesterol, a crucial lipid molecule with diverse cellular functions. Out of the 28 upregulated genes tested, three genes (*Mvk, Pmvk,* and *Idi1*) were found to be associated with this pathway. The second significant pathway, the Metabolism of steroids (R-MMU-8957322), involves various processes related to the metabolism of steroid molecules. Similarly, three genes (*Mvk, Pmvk*, and *Idi1*) were identified as associated with this pathway.

The REACTOME analysis using downregulated genes identified significantly enriched pathways. Among them, the Xenobiotics pathway (R-MMU-211981) showed enrichment, indicating the potential involvement of *Cyp3a11, Cyp3a59,* and *Cyp3a25* genes. Additionally, the Attenuation phase pathway (R-MMU-3371568) exhibited enrichment, with *Hspa1b* and *Hspa8* genes among the upregulated genes. Furthermore, the pathway Cytochrome P450 - arranged by substrate type (R-MMU-211897) demonstrated enrichment, suggesting potential dysregulation of *Cyp3a11, Cyp3a59,* and *Cyp3a25*. Pathways associated with apoptosis, such as Caspase activation via Death Receptors in the presence of ligand (R-MMU-140534) and Caspase activation via extrinsic apoptotic signalling pathway (R-MMU-5357769), were also enriched, with *Ly96* and *Casp8* among the upregulated genes. Additionally, the pathway Scavenging of heme from plasma (R-MMU-2168880) and Binding and Uptake of Ligands by Scavenger Receptors (R-MMU-2173782) showed enrichment, suggesting potential dysregulation of *Cd163, Apol9a*, and *Hspa1b/Hspa8*, respectively.

## DISCUSSION

Variants in *EFCAB4B* gene have been implicated in the development of NAFLD and variants in the proteins CRACR2A and Rab46 play roles in inflammation and in diseases enhanced by inflammation. Since we observed that mice lacking the *Efcab4b* gene appeared phenotypically normal except the size and weight of the livers and the epididymal fat pads, we hypothesized that *Efcab4b* deficiency could impact the expression of genes important for liver development and function. In this study, we examined the hepatic transcriptome of *Efcab4b* deficient mice and identified biological functions and genes associated with hepatotoxicity, lipid metabolism, and metabolic disorders. In particularly, we demonstrated roles for DEGS in liver and bile development which could give an insight in to the large livers observed in the *Efcab4b^-/-^ mice*.

### Liver disease related genes

To explore if proteins translated from the DEGs observed in *Efcab4b^-/-^* liver tissue could play a role in the progression of liver disease, we considered the molecular pathways that promote the progression of a healthy liver to a cirrhotic liver, and which pathways DEGs have been previously shown to have a role in (Figure 8). A simple NAFLD/ NASH-based literature search revealed associations between 32 DEGs and NAFLD/ NASH/ HCC (Tables 3 and 4) and the potential role they play in NAFLD progression (Figure 8).

**Figure 7.**
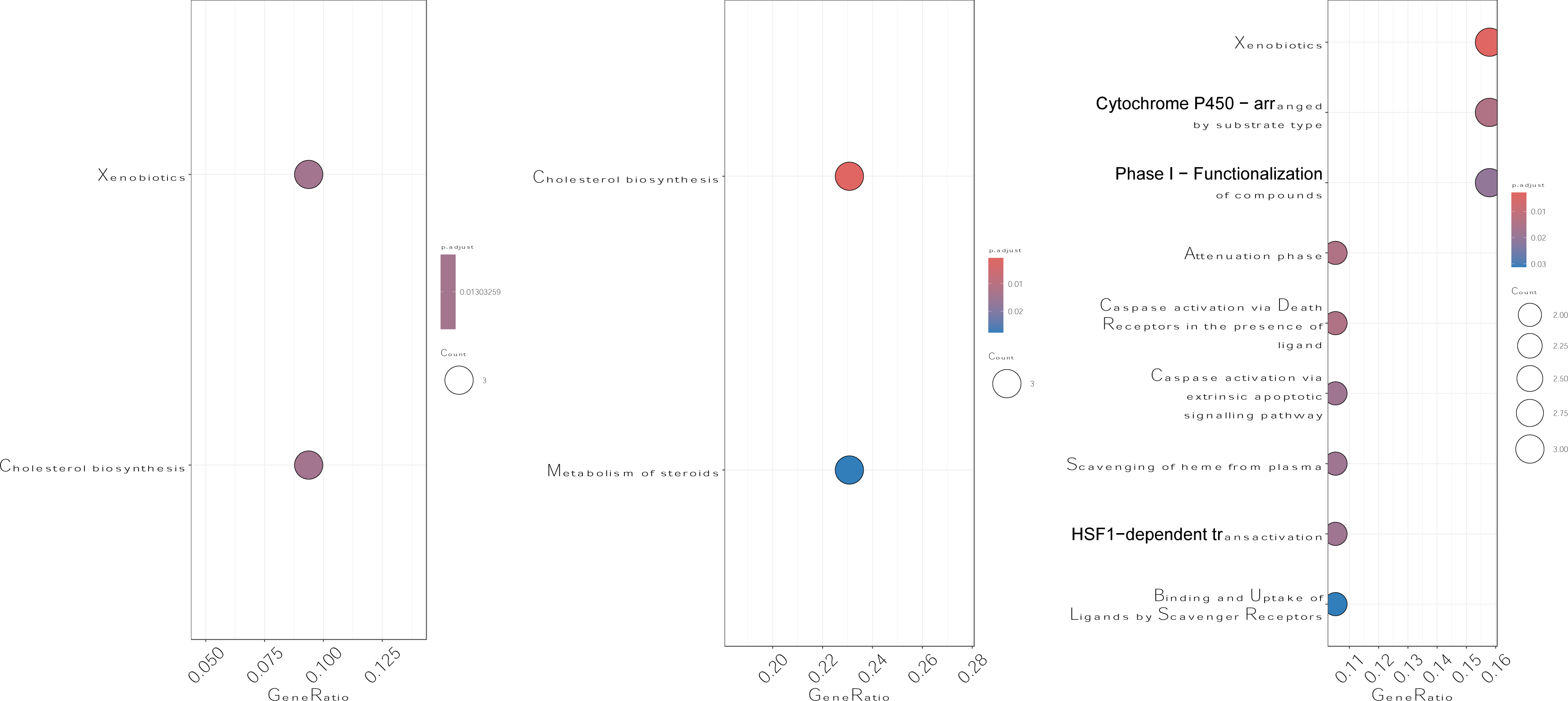
REACTOME analysis of all differentially expressed genes. (A) REACTOME analysis of all DEGS; (B) REACTOME analysis of upregulated DEGs; (C) REACTOME analysis of downregulated DEGs. The size of the circle represents the number of genes enriched in the pathway and the colour of the circle indicates the level of significance (P<0.05).

**Figure 8.**
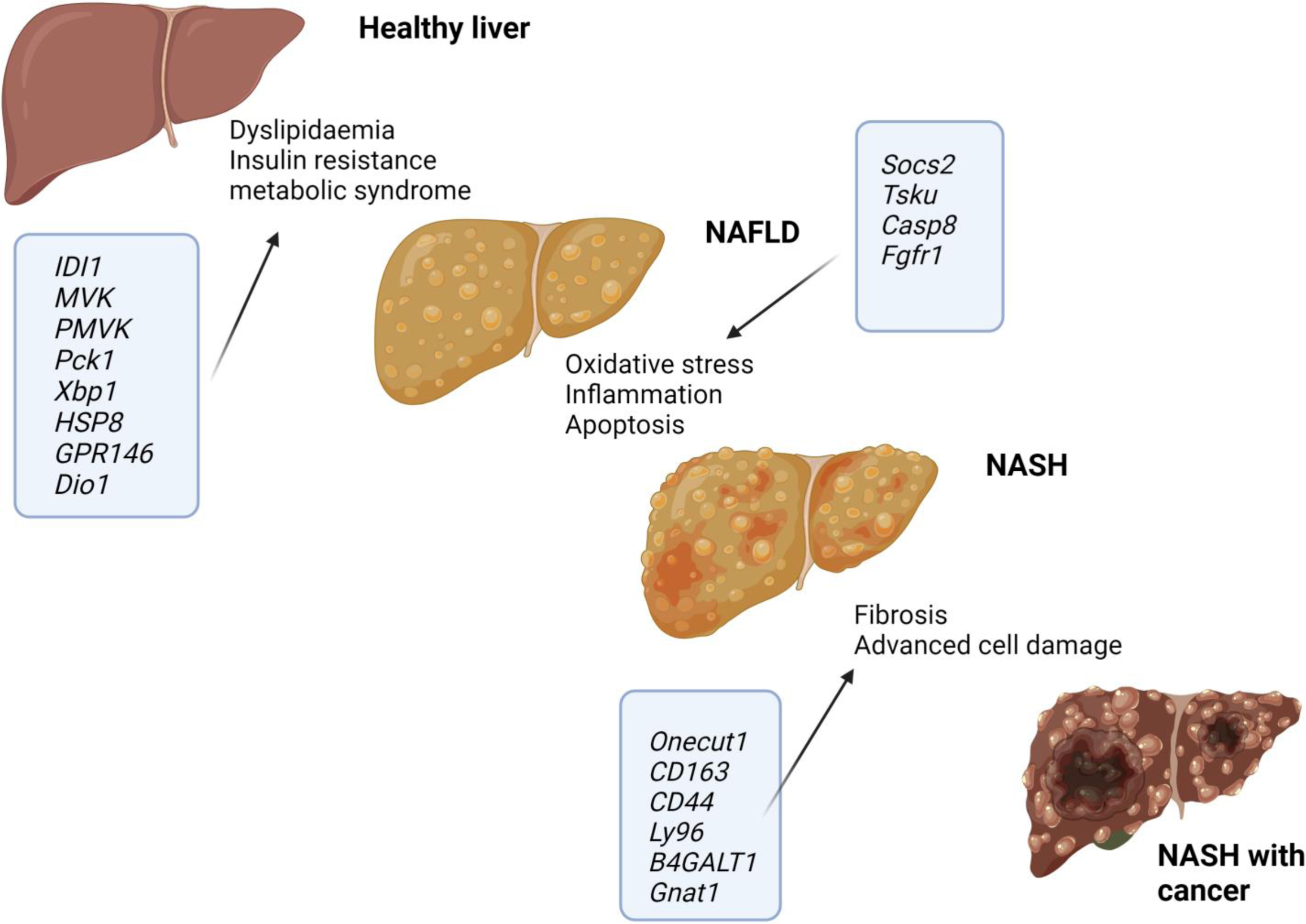
Schematic depicting the differentially expressed genes associated with molecular pathways that evoke the progression of NAFLD to NASH and HCC.

### Lipid metabolism and steatosis

Several identified DEGs suggest an association between *Efcab4b^-/-^* and the first stage of NAFLD: that is simple steatosis where the accumulation of lipid droplets within hepatocytes is associated with negligible, if any, inflammation. Several DEGs play roles in lipid metabolism and therefore have the potential to promote dyslipidaemias. For example, expression of inhibitor of DNA binding1 (*ID1*), phosphomevalonate kinase (*PMVK*), G protein-coupled receptor 146 (*GPR146*) ^33^ and X-box binding protein-1 (*XBP1*) are upregulated and phosphoenolpyruvate carboxykinase 1 (*PCK1*) downregulated in NAFLD or lipid disorders, this is reflected in the livers of *Efcab4b^-/-^* mice as compared to WT. *Efcab4b* depletion induced 1.488x downregulation in the expression of *Pck1*. *PCK1* is a gene that plays a critical role in hepatic glucose metabolism. It is involved in gluconeogenesis, the process by which the liver produces glucose from non-carbohydrate precursors, by catalyzing the conversion of oxaloacetate to phosphoenolpyruvate, a key step in glucose synthesis. Murine depletion of *Pck1* promotes metabolic-associated fatty liver disease (MAFLD) ^34^ and *PCK1* is down regulated in hepatocytes extracted from human diseased liver ^35^. Moreover, dietary plant sterols (phytosterols), that have cholesterol-lowering properties and attenuate deleterious effects of cholesterol overload, can rescue the impact of an HFD in a hamster model by restoring hepatic *Pck1* expression ^36^. *ID1* and *PMVK* genes encode for enzymes that are crucial for the cholesterol biosynthesis pathway and these genes are upregulated in the *Efcab4b^-/-^*mice versus WT mice (Table 3). Hepatic free cholesterol overload results in cholesterol-associated steatohepatitis and may play an important role in the development and progression of NAFLD, NASH and hepatic cancer ^37^. Statins appear to provide significant benefit in preventing progression to NASH and NASH-cirrhosis, suggesting in addition to cholesterol in the diet, the biosynthesis pathway plays a role. Betaine, a drug known to effectively improve hepatic lipid metabolism, evokes depletion of *IDI1* ^38^, whilst gypenosides, natural drugs used to treat lipid disorders, reduce the expression of *PMVK* ^39^. The upregulation of the transcription factor *Xbp1*, in the *Efcab4b^-/-^* mice has also been demonstrated in samples from patients with NAFLD, NASH and HCC ^40^. Hepatocyte-specific *Xbp1* deficiency inhibited the development of steatohepatitis in mice fed the high-fat diet whilst macrophage-specific *Xbp1* knockout mice developed less severe steatohepatitis and fibrosis than wild-type *Xbp1* mice. *XBP1* is the key transcription factor for initiating the unfolded protein response (UPR) in response to ER stress. In addition to being required for de novo fatty acid synthesis in the liver (a function unrelated to its role in the UPR ^41^), upon ER stress *XBP1* specifically induces expression of the transcription factor *FOXA3* and exacerbates lipid accumulation, linking ER stress to NAFLD progression ^42^. The analogous changes of these genes in *Efcab4b^-/-^*mice suggest the depletion of *Efcab4b* could evoke NAFLD progression by impacting lipid homeostasis. However, some DEGs that are positively associated with hepatic steatosis such as deiodinase 1 (*Dio1*) ^43^, are downregulated in the *Efcab4b^-/-^* livers, in addition the histological staining of the livers from *Efcab4b^-/-^*or WT mice did not show any pronounced lipid accumulation (in this particular study). Thereby we questioned if the identified DEGs could play a role in the progression of NAFLD from a simple macrovesicular steatosis to NASH and eventually hepatocarcinoma (HCC).

### Inflammation and ER stress in NASH

Progression from NAFLD to NASH involves other damaging factors, such as activation of inflammatory pathways, ER stress and dysregulated hepatocyte apoptosis. Several of the DEGs identified in the liver of *Efcab4b^-/-^* mice could play roles in these processes, however in these instances the expression in *Efcab4b^-/-^*liver contrasts that of disease. For example, in NASH, suppressor of cytokine signaling 2 (*SOCS2*) is down regulated whilst Tsukushi (*TSKU*), and Caspase 8 (*CASP8)* are upregulated. The contrasting expression levels of these genes in the livers of *Efcab4b^-/-^* mice suggest that the deletion of *Efcab4b* could be protective against inflammation. *Socs2* displays the highest fold upregulation (2.24x) in *Efcab4b^-/-^*mice. *SOCS2* is one of classic negative regulators of cytokine signaling, which has recently been described as an anti-inflammatory mediator. In human samples the level of *SOCS2* expression was negatively correlated with NASH: *SOCS2* overexpression in macrophages suppressed inflammation and apoptosis via inhibiting NF-κB signaling pathway, whilst *SOCS2* knock-down in macrophages caused an increased activation of NF-κB. In addition, *SOCS2* expression in macrophages also suppressed inflammation via limiting the activation of inflammasomes, strongly indicating that *SOCS2* plays a role in inhibiting inflammation and apoptosis via NF-κB and inflammasome signaling pathway in macrophages during NASH ^44^. Similarly, in HCC samples, immunohistochemical staining demonstrated lower levels of *SOCS2* protein expression in patients with HCC ^45^, whilst Liu *et al* suggested *SOCS2* is a protective factor, because its high expression improves the prognosis of HCC patients ^46^. Caspase 8 (*CASP8*) is essential for death-receptor-mediated apoptosis activity apoptosis, a critical mechanism contributing to inflammation and fibrogenesis, therefore its modulation is critical for the pathogenesis of NASH. Surprisingly the *Efcab4b^-/-^* mouse showed a 0.71x downregulation of *Casp8* expression which is protective in the development of NASH in *Casp8* knockout mice. Here, the lack of *Casp8* expression in hepatocytes reduced the diet-dependent increase in apoptosis and decreased expression of proinflammatory cytokines as well as hepatic infiltration. As a consequence, ROS production was lower, leading to a reduction in the progression of liver fibrosis in *Casp8^-/-^* livers ^47^. In agreement, curcumin treatment was shown to be beneficial in preventing the development of NASH in rat models by reducing apoptosis and decreasing the expression of *Casp8* ^48^. *TSK* is a hepatokine induced in response to both endoplasmic reticulum stress and inflammation in severely obese mice. In humans, hepatic TSK expression and increased serum levels are also associated with steatosis and acute liver failure ^49^. TSK level is downregulated by 1.07x in the livers from *Efcab4b^-/-^* mice versus WT mice.

### Fibrosis and HCC

Our RNAseq analysis of the livers from *Efcab4b^-/-^* mice exhibit downregulation of the macrophage marker CD163, the CD44 antigen and Lymphocyte antigen 96 (*Ly96*) and upregulation of beta-1, 4-galactosyltransferase 1 (*B4GALT1*). In previous studies increased circulating levels of CD163 in patients with HCC and in diabetic patients with advanced NASH fibrosis, suggests CD163 as a biomarker for disease severity in NAFLD^50^ ^51^. CD44 is a cell surface antigen that acts as a co-receptor in tyrosine kinase receptor signaling and is thereby involved in cell-cell interactions cell adhesion and migration. It is highly expressed in cancer cells and promotes tumour progression. Accordingly the upregulation of CD44 is observed in HFD diet fed animals displaying inflammation, fibrosis and is particularly significantly associated with the malignant transformation of hepatocytes in NAFLD ^52^ ^53^. The deletion of *CD44* inhibits metastasis formation in mice ^54^ and in obese patients, hepatic CD44 and serum sCD44 strongly correlated with NASH ^55^. Indeed, CD44 is considered a cancer stem cell marker ^56^ where immunohistochemical analysis of human HCC samples demonstrate increased expression of CD44. Ly96 (also known as myeloid differentiation factor 2: MD2) is a co-receptor for Toll-like receptor 4 (TLR4). Together they are key in recognition of lipopolysaccharide (LPS) and activation of proinflammatory pathways. In a mouse model of NASH, knockout of *Ly96* significantly attenuated triglyceride accumulation, lipid peroxidation, inflammation and liver fibrosis ^57^. Similarly an angiotensin II liver injury mouse model displayed significantly reduced inflammation and fibrosis ^58^. *B4GALT1* is significantly down regulated in the livers of patients with HCC and reducing *B4GALT1* enhanced HCC cell migration and invasion in vitro and promoted lung metastasis of HCC in NOD/SCID mice ^59^.

Comparing the level of gene expression (i.e. those that are upregulated in disease (yellow) or down regulated in disease (blue)) between previous NAFLD/ NASH/ HCC studies (Tables 3 and 4), to the expression levels in the livers of the *Efcab4b^-/-^* mouse, demonstrates that of the 33 genes, 14 genes are expressed in a similar manner in the *Efcab4b^-/-^*mice, whilst 19 are contradictory. Particularly, DEGs that have been shown to have roles in molecular pathways leading to NASH, through inflammation, are expressed in a contradictory manner suggesting depletion of *Efcab4b* could be protective and previous studies suggest a role for Rab46 in inflammation. However, the interpretation of this data is limited to the model being a global depletion of *Efcab4b* and thereby the impact on gene expression in the liver tissue may not be direct. Moreover, depletion of *Efcab4b* prevents expression of both CRACR2A and Rab46 protein isoforms. Since the SNP that has been associated with NAFLD could play a role in splicing it may be that the balance of expression between these 2 proteins has an impact. Functional cellular assays and rescue of isoform expression would lead to a greater understanding of the mechanisms by which *Efcab4b* impacts liver function in health and disease.

## CONCLUSION

In summary, this discovery may uncover the intricate regulatory mechanisms underlying the CRACRR2A/ Rab46 phenotype and provides valuable insights into the functional implications of these genes. Our findings may contribute to a better understanding of the molecular landscape associated with Rab46, opening avenues for further investigation into the biological processes and potential therapeutic targets associated with non-alcoholic fatty liver disease.

## Author Contributions

CWC performed genetic analysis. LP undertook the mouse studies. LP, CMA, FD, JS, NF and LM obtained the liver samples. CWC, LP, LM analysed the data. LM conceived the study and co-wrote the article with CWC and reviewing from all authors. LM generated research funds and coordinated the project. All authors have read and agreed to the published version of the manuscript.

## Acknowledgments

The study involved use of ARC4, which is part of the High Performance Computing Facility at the University of Leeds. We’d like to thank LeedsOmics for all their input (https://omics.leeds.ac.uk/).

